# Altered sperm DNA methylation in overweight men associates transposable element regulation to paternal origins of disease risk in children

**DOI:** 10.1101/2025.09.15.675616

**Authors:** Mathieu Schulz, Xiaojian Shao, Romain Lambrot, Christine Lafleur, Donovan Chan, Marie-Charlotte Dumargne, Amanda J. Macfarlane, Sergey Moskovtsev, Guillaume Bourque, Tomi Pastinen, Clifford Librach, Hope Weiler, Jacquetta Trasler, Sarah Kimmins

## Abstract

The occurrence of childhood neurodevelopmental disorders has been steadily increasing for decades yet we have little understanding of modes of inheritance implicated in these diseases. Epidemiology studies have revealed an association between an elevated paternal BMI and an increased risk for autism in children, suggesting non-genetic modes of inheritance may be involved. Epigenetic marks, like DNA methylation at cytosines, are especially susceptible to environmental, diet and lifestyle changes. Yet, whether a man’s BMI influences the sperm methylome and potentially impacts offspring development remains unresolved. Using MethylC capture (MCC)-sequencing, we identified over 38 000 differentially methylated CpGs (DMCs) in the sperm of men with an elevated BMI, with many occurring in regions that were enriched for neural gene and disease ontologies. Differentially methylated regions (DMRs) were enriched at transposable elements that can act as active enhancers during human zygotic genome activation, and at gene promoters for early lineage specification. These results suggest that sperm methylome alterations that are linked to an elevated BMI may influence key embryonic transcriptional process, notably those associated with trophectoderm and placental development.

## Introduction

Epigenetic modifications are dynamic gene regulators that are shaped by environmental influences ^1^. DNA methylation, which consists of cytosine methylation in CpG dinucleotides, is associated with gene repression, playing a pivotal role in regulating gametogenesis and early embryonic development. In mice, DNA methylation is a central regulator of spermatogenesis, male fertility and early embryonic development. Namely, DNA-methylated imprinted genes, characterized by their monoallelic and parent-of-origin expression, have an extraordinary ability to resist the epigenetic reprogramming that occurs post-fertilization in the early mammalian embryo ^2^. These genes have an unequal effect on embryonic development, with the paternal genome promoting the formation of extra-embryonic tissues ^3,4^. Along with the establishment of paternally methylated regions, DNA methylation at discrete regions serves essential roles in spermatogenesis, including silencing of transposons or regulation of gene expression ^5–7^. Moreover, the transmission of the paternal epigenome at fertilization is important for zygotic genome activation ^8,9^. It is therefore plausible that errors in paternally inherited DNA methylation could alter embryo transcription programs and lead to higher risk for disease phenotypes in children.

Indeed, abnormalities in the establishment of DNA methylation in the developing germline have been associated with epigenetic inheritance between generations, mostly through imprinted genes. For example, Prader-Willi syndrome, a complex disorder characterized by hyperphagia and problems in neurodevelopment can be attributed to abnormal imprinting near the paternally expressed gene *Small Nuclear Ribonucleoprotein Polypeptide* (*SNRPN*) ^10^. Similarly, differential sperm DNA methylation in the *SNORD* gene family found within the Prader-Willi gene cluster is associated with an increased risk of autism in children ^11^. In fact, and despite high variabilities across continents and regions, a global rise in the rates of autism can be observed worldwide (i.e., 63.7% increase in 2016 vs 2008 in the USA) ^12^. Notably, recent work estimates that prevalence of autism in 8 year olds across the United States is 1 in 36 ^13^. These staggering increases also coincide with rising rates of paternal obesity and the societal trend of delaying parenthood. Notably, paternal obesity increases the risk of having a child with autism (odds ratio of 1.73) ^14^, as does advanced paternal age (55% increased risk) ^15,16^. Taken together these findings suggest that advanced paternal age, metabolic health, and being overweight may be implicated in the rising rates of neurodevelopmental disorders in children. How these paternal factors affect the sperm methylome and in turn the next generation remains poorly understood ^17^.

Indeed, while epidemiological studies support the notion of epigenetic inheritance, there are yet only a handful of studies that have tracked changes in the sperm epigenetic content to the embryo or children. Recently we demonstrated that the sperm DNA methylome is perturbed in association with altered folate metabolism or when exposed to the environmental toxicant dichlorodiphenyltrichloroethane (DDT) ^19,20^. Another recent study showed that paternal exposure to air pollution also alters the sperm methylome and is associated with lower birth weight ^21^. The mechanism by which the paternal sperm methylome could exert its impact in early development represents an active field of research. One route of transmission may involve transposable elements. Indeed, DNA methylation is a well-studied silencer of transposable element activity in the germline, which directly affects fertility and early human development ^22^. Namely, the silencing of LTR-transposable elements in early human development is critical for zygotic genome activation ^23^. These elements are particularly interesting candidates for bridging generations epigenetically, some are known to evade the global DNA methylation erasure that rewires the methylome of the embryo in its first days of development ^24,25^. Moreover, they constitute central components of gene regulatory networks as they bear diverse transcription factor binding motifs, with some elements having evolved specific regulatory functions for the emergence of cell lineage in humans ^26–28^.

Despite advancements in sequencing, studies of the sperm methylome have mostly focused on gene promoters and imprinted regions, overlooking the dynamic and underexplored intergenic regions that comprise a significant portion of the genome. Additionally, the potential link between sperm methylation patterns and early human embryonic development has been largely understudied. Addressing these gaps is essential toward uncovering the mechanisms underlying the effects of paternal health on offspring development.

To investigate the association between paternal BMI and sperm DNA methylation, we analyzed sperm from men with BMIs ranging from normal to overweight and obese using MethylC Capture (MCC)-seq, a high-resolution method targeting over three million CpG sites, including environmentally sensitive regions ^19^. Specifically, this panel interrogates over three million CpGs in the genome, including the > 850,000 sites present on the Infinium MethylationEPIC BeadChip, an array-based technique targeting commonly assessed gene promoter, CpG island regions and enhancers, widely used in DNA methylation analyses. In addition, the human sperm MCC-seq panel targets approximately one million CpGs with “dynamic” levels of DNA methylation. These are environmentally sensitive sequences which demonstrate intermediate DNA methylation levels (typically between 20 – 80%) mostly found at intergenic regions. Thus, using this approach we gained insight into underexplored regions of the sperm methylome, while retaining our ability to compare our epigenetic signature with previously published datasets. In this study, our analysis identified over 38,000 differentially methylated cytosines (DMCs) in sperm from men with elevated BMI, that showed a trend for hypermethylation. These DMCs were significantly enriched in intergenic regions, highlighting the importance of these often-overlooked regions. Gene ontology analysis revealed that many of these regions were associated with neural development, suggesting potential impacts on offspring brain function and developmental processes.

## Methods

### Ethical statements

This study was approved by the Faculty of Medicine Institutional Review Board, McGill University, the Health Sciences Research Ethics Board(A09-M57-15B), University of Toronto (IRB No. 30089) and the Health Canada-Public Health Agency of Canada Research Ethics Board (REB2019-0006).

### Participants and sample collection

In a prospective study we recruited 71 patients from the CReATe Fertility Center (Toronto, Ontario, Canada) from January 2016 to December 2017. Semen and blood samples were collected on the same day after men abstained from sexual activities for at least 2 days. Two aliquots of undiluted raw semen were collected 30 minutes after liquefaction. Semen and centrifuged blood samples were stored in a −80°C freezer for long-term storage and later analysis. Participants who reported use of tobacco products, recreational drugs, steroids, or had the common polymorphism 677C→T Methylenetetrahydrofolate reductase (*MTHFR*) ^29^ were excluded from the study.

We selected 52 samples for downstream analysis based on the participant having normal semen parameters according to WHO standards ^30^. Body Mass Index (BMI) was calculated from self-reported weight (kg)/ height^2^ (m), and participants were classified by BMI as normal (NOR =20, 18.5–24.9 kg/m²), overweight (OW =20, 25–29.9 kg/m²), or obese (OB =12, ≥30 kg/m²) following WHO criteria ^31^. We also obtained clinical data including fertility treatment and outcome (**Supplementary Table 2**). However, without a fertile sperm donor control group for comparison, a meaningful analysis relating sperm DNA methylation to clinical outcomes was not possible.

### Analysis of plasma and serum samples

We assessed plasma and serum parameters including serum folate (nmol/L), B12 (pmol/L), homocysteine (µmol/L), and 25-hydroxyvitamin D (25-OHD) (nmol/L). Serum and RBC total folate, plasma homocysteine, and serum vitamin B12 were measured according to the manufacturer’s instructions (Siemens ADVIA Centaur XP) at the Nutrition Research Canadian Health Measures Survey Reference Laboratory. Policies, procedures, and processes within the laboratory are aligned with licensing and accreditation standards of clinical laboratories in Canada and specifically the Institute for Quality Management in Healthcare, and assay performance targets (e.g., precision, bias, specificity, sensitivity) reflect their recommendations. All assays were verified prior to sample analysis using third-party quality control materials (BioRad) ^32^. Red Blood Cell (RBC) samples were diluted 1:21 with ascorbic acid solution for folate analysis. After being mixed, hemolysates were allowed to stand for 45 min at room temperature. RBC folate levels were then normalized to measured hematocrit. RBC total folate concentrations that fell above the highest calibrator were assigned the value of the highest calibrator for analyses. Serum 25(OH)D concentration was measured using the LIAISON Total 25(OH)D assay (DiaSorin Inc, Stillwater, MN, USA) at McGill University in the School of Human Nutrition. The intra-assay and inter-assay CVs were <6.5%; the accuracy was 95-106% for NIST972a Level 1 (Vitamin D Metabolites in Frozen Human Serum) and 91-101% for the DiaSorin 25OHD Controls using the mid-range of manufacture specifications. Values of 25(OH)D were standardized as recommended by the Vitamin D Standardization Program (Durazo-Arvizu et al., 2017).

### Sperm DNA isolation

Sperm genomic DNA extraction was performed as described in ^20^. Briefly, 10 million spermatozoa were lysed overnight at 37°C in a buffer containing a final concentration of 150 mM Tris, 10 mM ethylenediaminetetraacetic acid, 40 mM dithiothreitol, 2 mg/mL proteinase K, and 0.1% sarkosyl detergent. DNA was then extracted using the QIAamp DNA Mini kit (Qiagen) according to the manufacturer’s protocols. Next, the purity of the extraction was checked using bisulfite pyrosequencing against imprinted loci to screen for somatic cell contamination. Briefly, 500ng of sperm DNA was used for bisulfite conversion with EpiTect bisulfite kit (Qiagen) following the manufacturer’s instructions. Then, regions of interest (paternally and maternally methylated ICRs of imprinted genes) were amplified by PCR using biotinylated primers ^29^ (**Supplementary Table 2**). Biotinylated strands were captured using streptavidin-coated Sepharose beads before being washed. Then, pyrosequencing reaction was performed on a PyroMark Q24 workstation (Qiagen), using the PyroMark Q24 kit (Qiagen) following the manufacturer’s instructions. Following this pyrosequencing reaction, the exclusion criteria for somatic contamination of sperm DNA preparation were determined at methylation levels below 90% ICRs of paternally methylated imprinted gene (*H19*) and maternally methylated imprinted genes (*MEST*, *SNRPN*). All samples in the current study demonstrated no somatic cell contamination (**Extended Data Fig.1**, **Supplementary Table 2**).

### Sperm DNA methylation Capture (MCC-seq)

MCC-seq sequencing was performed as described in ^19,20,33^. Briefly, 1 – 2μg of DNA were sonicated (Covaris) and fragment sizes (300 – 400 bp) were controlled on a Bioanalyzer DNA 1000 Chip (Agilent). Then we performed library preparation, following the KAPA Biosystems’ protocols for DNA end repair, 3′-end adenylation, adaptor ligation, and clean-up steps. Using the Epitect Fast bisulfilte kit (Qiagen), samples were bisulfite converted according to the manufacturer’s protocol, followed by quantification with OliGreen (Life Technology). Library amplification was performed with PCR (9 – 12 cycles) using the KAPA HiFi HotStart Uracil + DNA Polymerase (Roche/KAPA Biosystems). Final WGBS libraries clean-up was made with Agencourt XP AMPure beads (Beckman Coulter), validated on Bioanalyzer High Sensitivity DNA Chips (Agilent), and finally quantified using PicoGreen (ThermoFisher). Following WGBS library preparations of all samples, CpGs targeted by our recently developed human sperm panel ^19^, were captured using the SeqCap Epi Enrichment System protocol by RocheNimbleGen. Our custom SeqCap Epi panel covers 3,179,096 CpG sites which are found mainly in intergenic (35%), intronic (33%), and promoter-TSS (19%) regions. Of the captured CpGs, 937,141 have been selected for inclusion in the capture panel based on demonstrated environmental sensitivity and high variability, with DNA methylation levels ranging from 20 – 80% in a WGBS dataset of pooled sperm from 30 men. These were termed “Dynamic” CpGs (Chan et al., 2019). Other selected CpGs corresponded to sites from the commonly used Infinium MethylationEPIC BeadChip array which targets gene promoters / CpG island regions and enhancers ^19^. Equal amounts of multiplexed libraries (84 ng of each; 12 samples per capture) were combined to obtain 1 μg of total input library and was hybridized at 47°C for 72h to the capture panel, specifically the human sperm capture panel recently developed by our group. This was followed by washing, recovery, PCR amplification of the captured libraries and final purification, according to manufacturer’s recommendations. Quality, concentration, and size distribution of the final captured libraries were determined using Bioanalyzer High Sensitivity DNA Chips (Agilent). The capture libraries were sequenced with a 200-cycle S2 kit (100 base paired-end sequencing) on the NovaSeq 6000 following the NovaSeq XP workflow.

### MCC-seq data pre-processing

The targeted MCC-seq data of sperm were processed using the Genpipes pipeline ^34^. Specifically, the high-throughput bisulfite sequencing reads (i.e. paired-end raw FASTQ reads) of MCC-seq were first trimmed using Trimmomatic (version 0.36) for quality (phred33 >= 30) and short reads (length n > 50 bp), as well as Illumina adapters ^35^. The trimmed reads were then aligned to the *in silico* bisulfite-converted human reference genome (hg19 / GRCh37 version) using Bismark (version 0.18.1) ^36^. Particularly, the alignments were performed with the bowtie2 aligner (version 318 2.3.1) in paired-end mode, using the non-directional protocol setting and other default parameters ^37^. The aligned bam files per lane were merged and then de-duplicated using Picard (version 2.9.0) ^38^. Subsequently, cytosine-level methylation calls were obtained using Bismark, which recorded counts of methylated and unmethylated reads at each cytosine position in the genome. DNA methylation level of each CpG was defined as the number of methylated reads divided by the total number of sequenced reads at that CpG. Additionally, CpG sites overlapping with SNPs (dbSNP 137), the DAC Blacklisted Regions or Duke Excluded Regions (generated by the Encyclopedia of DNA elements - ENCODE project) ^39^ or were removed. CpG sites with less than 20X read coverage were also discarded.

### Single nucleotide polymorphism and genotype analysis

From the deduplicated bam files, variants (including homozygous alternate and heterozygous genotypes) were inferred using BiSNP (version 0.82.2) ^40^. A set of unique variants called from all individuals was used as the potential SNP set. The homozygous reference genotypes of individuals on these SNPs were extracted from the aligned bam files, requiring a read coverage of no less than 10X aligned to the reference allele. Principal component analysis (PCA) was conducted on SNPs with genotypes inferred from all individuals to assess the genetic variations across the group.

### Differentially methylated CpGs (DMCs) and regions (DMRs) identification

Generalized linear regression models (GLMs) were constructed using the methylation proportion, inferred from the combination of methylated reads and unmethylated reads, as a binomially distributed response variable to investigate associations between DNAme in sperm and BMI. Both continuous and binary models were explored, and the models were adjusted for age and other serum covariates including RBC folate, Serum folate, Serum B12, Homocysteine, 25-OHD. For the downstream analyses, used continuous regression. For certain CpGs, used a cut-off for sequencing coverage (>= 20X), and any CpGs below this threshold were removed from our analysis. Additionally, non-variable CpGs (standard deviation = 0) were removed to reduce the multiple testing burden. R function glm(), with the binomial family were used to fit each model, and p-values for variables of interest were obtained accordingly. The obtained p-values were then corrected by estimating the false discovery rate q-values using the Bioconductor/R package qvalue (version 2.16) ^41^. We defined significant associated DMCs when q-values were less than 0.01. Furthermore, consecutive DMCs (at least 3) spanning 500 bp were merged to define differentially methylated regions (DMRs). For the statistical analysis, R version 3.6.0 was used along with Bioconductor packages from the 3.9 release.

### Enrichment and Gene ontology analyses

Genomic features and Repeat masker annotations were retrieved using AnnotationHub (version 3.8.0) and annotatr (version 1.26.0) packages from Hg19 assembly. To determine the association between regions set and features of interest, Zcores for enrichment of DMCs over genomic features and transposable elements were calculated by using permTest() function from RegioneR (version 1.32.0) using 1000 permutations, masking for Hg19 blacklisted regions ^39^. As a control, we used the resampleRegion function to randomize a set of regions issued from the background list of MCC-seq covered CpGs.

To determine Gene/Disease Ontologies associations, the overlap of DMCs and DMRs at genic features (including promoters, exons and introns) were retrieved using IRanges (version 2.36.0). Then we used ClusterProfiler (version 4.8.3) ^42^ to estimate functional associations using Fisher exact Test, with p-values adjusted with the Benjamini-Hochberg (BH) method. Cut-offs for p-value and qvalue were set at 0.05 or below, only top enriched ontologies were displayed, with no additional arbitrary filtering performed.

### Integration of publicly available datasets

To gain deeper biological context, we leveraged publicly available human datasets. First we overlapped our BMI associated sperm DMRs with our previously generated H3K4me3 human sperm peaks ^43^. We also compared with sperm DMRs associated with paternal age ^33,44^. Then, to gain more insight into early human development, we sought to use 8 cell-stage and blastocyst stage embryo enhancer chromatin landscape (H3K4me3, H3K27ac and ATAC) and gene expression datasets ^45–47^. Enhancers at the 8 cell stage were defined based on the overlap of H3K27ac and/or ATAC peaks and their association with gene promoters, as identified through HiC ^47^. Additionally, a maximum distance threshold of 200000bp was applied between 8C-Enhancer and their associated genes. For transcriptomic public datasets, count per million (CPM) normalization was conducted using EdgeR (version 3.42.4), and differential gene expression analysis was carried out using voom transformation from the limma R package (version 3.56.2). PCA was performed considering all detected transcripts. P-values were determined using limma and adjusted with BH correction. The threshold for differentially expressed genes (DEGs) was set to a false discovery rate (FDR) < 5% and at least |log2 (fold change)|> 1.5. Overlaps between public datasets, genomic annotations and our sperm DMCs/DMRs were identified using the IRanges package (version 2.36.0).

### Transposable element insertion age analysis

To analyze how sperm DNA methylation changes are associated with specific transposable elements (TEs) according to their putative insertion age, we used the “percentage.divergence” from consensus motifs provided by Repeat masker as an “age”-proxy of each TE. Briefly, a lower divergence percentage is associated with a younger element. We then applied a linear regression model using the stats package (version 4.3.1), with changes in DNA methylation at sperm-DMCs overlapping with TEs as the response variable, and the divergence percentage of TEs as the predictor. To control for this model, we ensured homogeneity of predictions over changes, and homogeneity of variances using the performance package (version 0.11.0). From this model, we retrieved the R^2^ determination coefficient, Pearson coefficient as well as p-value calculated via F-test.

### Binding motifs analysis

To retrieve binding motifs of transcription factors at sperm DMRs overlapping with transposable elements, 8-cell stage enhancer and gene promoters, we used the monaLisa package (version 1.6.0). Briefly sperm-DMRs were binned according to their methylation changes (gained or lost methylation in participants with elevated BMI compared to NOR individuals). Binding motifs were retrieved form the JASPAR 2020 database ^48^. Enrichment of binding motifs was calculated over bins using Fisher Test and default monaLisa (version 1.6.0) parameters from the calcBinnedMotifEnrR() function ^49^. The analysis of retrieved binding motifs was performed using metascape ^50^ to extract transcription factor broad functions. We limited redundancy of pathways and ensured overall relevance with early embryonic development.

### Data Visualization

The R package ggplot2 (version 3.5.1) was mainly used for data visualization alongside with these packages:

- Manhattan plot was generated using karyoploteR (version 1.26.0) ^51^
- Bubble plots for Ontologies were made using ClusterProfiler (version 4.8.3) ^42^
- Area-Proportional Venn diagram was done using eulerr (version 7.0.2) ^52^
- Upset plot was generated using UpSetR (version 1.4.0) ^53^
- Tree plots and heatmaps of binding motifs were made using monaLisa (version 1.6.0) ^49^
- Tracks visualizations were generated using plotgardener (version 1.6.4) ^54^

## Results

### Men with an elevated BMI showed differential sperm methylation

Extensive epigenetic remodeling occurs in germ cell development *in utero* and continues in adulthood throughout spermatogenesis ^7,55^. Thus the window for environmental influence on the sperm methylome extends beyond pre-natal development as has been demonstrated in the case of exposure to endocrine disrupting chemicals ^20^ and folate supplementation ^19^. Notably an altered sperm methylome is also implicated in male infertility ^56^. To investigate whether there is an association between a man’s BMI and sperm DNA methylation, we collected sperm from men with normal semen parameters according to the WHO ^30^ as controls, and a range in BMI from overweight to obese (25 ≤ BMI and above). Clinical characteristics are detailed in **Figure 1A, B**. We integrated serum parameters for each individual to control for potential covariates. Notably, RBC folate range was adequate (661.50–1879.06 nmol/L, mean: 1182.54 nmol/L), along with serum folate (mean: 42.62 nmol/L), serum B12 (mean: 384.89 pmol/L), homocysteine (mean: 11.19 µmol/L) and 25-OHD (mean: 54.578 nmol/L). These methods indicate a mean BMI of 27.3 ± 4.5 kg/m² (**Fig. 1A**). While the average age of the patients is 37.3 years old, we observed no significant differences in the ages of individuals across different BMI groups, except for a slightly younger age in obese patients compared to overweight patients (p-value=0.03, **Figure 1B**). All parameters previously mentioned can be found in **Supplementary Table 1**.

**Fig 1.**
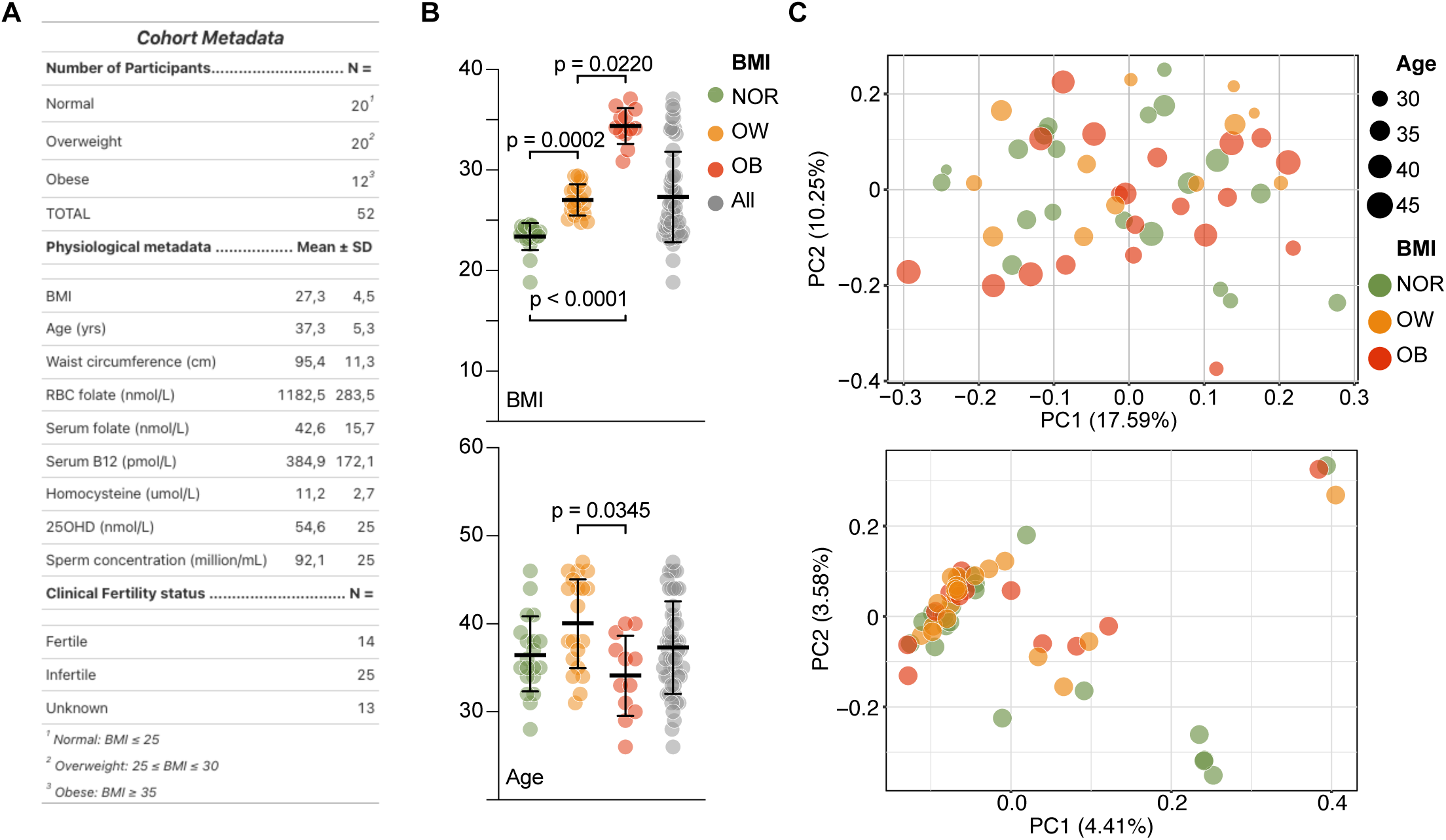
BMI contribute to variability observed in sperm methylome. **A**. Summary data of study Cohort. RBC Folate (red blood cell folate). 25OHD (25-hydroxy Vitamin D). All participants presented normal semen parameters according to WHO standards. **B**. Barplot of BMI (top) and Age (bottom) repartition across 3 groups of participants. Grouping was made according to BMI; Obese (OB) ≥35, Overweight (OW) 25≤BMI≤29.9 and Normal (NOR) BMI≤ 24.9. Data displayed is mean with SD, statistical test is ANOVA with Kruskall-Wallis. **C**. Principal Component analysis showing contribution to variability of BMI and Age (top) and Genotype (bottom) in top 1% differentially methylated cytosines.

To investigate the sperm DNA methylation landscape, we applied a customized MethylC-Capture followed by sequencing ^19,20^ (**Extended Data Fig. 1**). Overall, the average of read coverage of CpGs per sample is 30X. After filtering CpGs with low read coverage (<=20X), a total number of 2 404 135 CpGs covered by at least one sample were remained. To further remove CpGs with low sample coverage, we obtained 1,835,671 CpGs with 20X coverage and covered by more than 30 samples (i.e., 75.35% of total CpGs were retained after removing low sample coverage CpGs). Furthermore, 1 937 CpGs showed non-variable methylation (0.08% of total CpGs, or 0.11% after removing low sample coverage CpGs) (**Extended Data Fig. 1A, 1B, 1C**) ^19^.

First, epigenetic variation in global DNA methylation was assessed by Principal component analysis (PCA) to determine whether samples would be discriminated in relation to age, BMI and genotype. While the first two principal components accounted for 17.59% and 10.25% of the variance (PC1 and PC2, respectively) in our dataset, there was no clustering by BMI or age (**Fig. 1C**). Meanwhile, using inferred genotype data, our genetic PCA analysis revealed minimal variance among individuals (**Fig. 1C**), indicating that there are no substantial genetic differences in our cohort and that global genetic effects are unlikely to drive the observed DNA methylation dynamics.

To identify whether there were differentially methylated CpGs (DMCs) in sperm that were associated with increased BMI, a quantitative association between sperm DNA methylation and BMI was established using a generalized linear regression model adjusted for potential covariates (age, folate, B12, homocysteine, and 25-OHD). Methylation gains and losses at DMCs were defined as a CpG showing a significant (qvalue <0.01) increase or decrease in DNA methylation level associated with BMI.

We detected 38719 DMCs, with 88% gaining methylation (34,234 DMCs) and 12% losing methylation (4485 DMCs) (**Fig. 2A and Extended Data Fig. 2A, 2B, Supplementary Table 3**). Analysis of CpG enrichment over background MCC-seq panel background, revealed DMCs were primarily enriched within intergenic regions and displayed intermediate levels of DNA methylation (Zscore > 100 and DNA methylation level is, in average, 45% ±6%). Some DMCs occurred in “genic context” regions (13.5% on exons and around 5.8% considering both 5’UTR and 3’UTR) (**Fig. 2B, 2C, Extended Data Fig. 2C**). Genes of interest gaining methylation include Myelin Transcription Factor 1 Like (MYT1L), that has neuroendocrine functions and has been implicated in syndromic obesity, and neurodevelopmental disorders including autism spectrum disorder and schizophrenia ^57–59^ (**Fig. 2A**). Likewise, a member of the neuronal specific catenin family, CTNNA2, involved in complex cortical dysplasia, showed a loss of methylation in sperm from men with elevated BMI ^60^. While the density analysis suggested there might be an accumulation of DMCs at telomeric and sub-telomeric regions, a hot-spot analysis revealed no clear telomeric regional bias (**Fig. 2A, Extended Data Fig. 2D)**. Likewise, probing specific transposable element families that could accumulate DMCs at these regions revealed no clear overrepresentation (**Fig. Extended Data Fig. 2E**). Overall, our MCC-seq analysis reveal that men with an elevated BMI demonstrate a trend for sperm DNA methylation gains genome-wide, with DMCs mostly found close to genes with known association to neural disorders.

**Fig 2.**
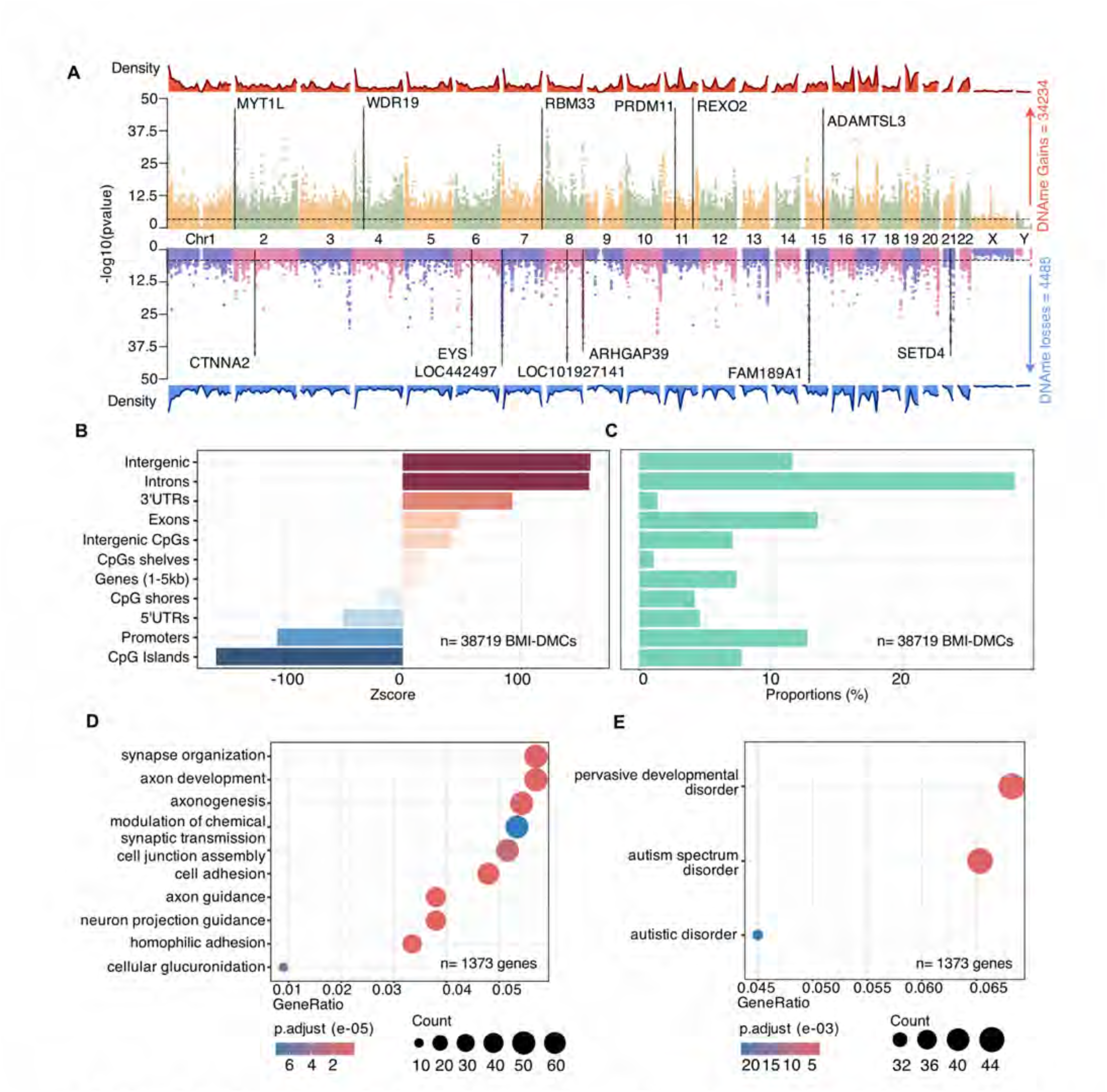
BMI mostly associate with hypermethylated DMCs at intergenic regions in the sperm and with neural gene ontologies and disorders. **A**. Manhattan plot showing the repartition of DNA methylation (DNAme) gains and losses in sperm BMI-DMC (n=38719) alongside chromosomes, according to their p-value (-log10). BMI associated DMCs were defined using General linearized Model, correcting for all covariates displayed in Fig1, and showing qvalue<0.01. Arrows associate genes bearing most significant hypo or hyper DMCs. **B**. Barplot of Zscore enrichment of BMI-DMCs (n=38719) on key genomic features over background. Scores were obtained using n=1000 random permutation t-test using R package RegioneR. **C**. Barplot of proportions of BMI-DMCs (n=38719) on key genomic features shown in percentage. **D, E**. Bubble plot of Gene/Disease ontologies enrichment analysis (R package ClusterProfiler) for BMI-DMRs located at gene promoters, introns or exons. Statistics are t-test for enrichment over background with BH correction. DMRs were defined using 500bp tile with 3 or more DMCs.

### Elevated BMI associated with DMCs found at neural gene and disease ontologies

To probe how differential DNA methylation may impact genome functionality, we merged differentially methylated CpGs into differentially methylated regions (DMRs). DMRs are more likely to exert regulatory functions than single DMCs. We observed over 3 828 sperm BMI-DMRs (**Supplementary Table 3**) among which, we found 1 373 unique genes bearing sperm BMI-DMRs (**Fig. 2D, 2E, Supplementary Table 4**). Using ClusterProfiler, we revealed the gene and disease ontologies most enriched at these DMRs were associated with neural development including autism spectrum disorder (**Fig.2D, E, Supplementary Table 5**). This observation resonates with epidemiological studies that showed a higher risk for children of fathers with and elevated BMI and neural disorders ^14,61^, although the underlying mechanisms and whether this involves epigenetic inheritance remains unclear.

Although epigenetic modifications in sperm are frequently studied alone, there are suggestions of functional interaction between DNA methylation and chromatin marks (notably histone modifications) in the establishment and role of the sperm epigenome across generations. In general, promoters with high CpG content contain higher levels of nucleosomes than CpG-poor intergenic and intronic space and are generally hypomethylated in sperm ^62,63^. However, breaking the general misconception DNA methylation and methylation at lysine 4 of histone 3 (H3K4ME) are mutually exclusive in sperm, over 24,000 H3K4me3 peaks overlap regions marked by intermediate (between 20 and 80%) and high (over 80%) DNA methylation ^43^ at developmental loci. Indeed, among our 3 818 BMI-DMRs in sperm, we observed 594 DMRs overlapping with a significant H3K4me3 peaks (log2 enrichment >3) (**Extended Data Fig.2F)**, most of them being significantly enriched at intronic (60%) and genic regions (13% at 1-5kb from TSS and exons) (**Extended Data Fig.2G, Supplementary Table 6**). Given the antagonism between DNA methylation and H3K4me3, and the strong association of the latter with active regulation of gene expression, this suggests that DMRs in the sperm of men with increased BMI could associate with gene mis-regulation in spermatogenesis, or even embryonic development. Such effects could be a direct influence on gene activation, or via changes in the epigenetic machinery.

### BMI impacts methylation at young LINE and LTR elements in human sperm

Transposable elements (TE) are drivers of genome organization, and act as regulators of gene expression. In spermatogenesis the disruption of DNA methylation at TE occurs in male infertility ^22,56^. In addition, the expression of specific transposable elements is required for early embryo development at zygotic genome activation (ZGA) ^23,47^, and lineage specification ^27,64^. Interestingly, a significant proportion of BMI-DMCs (n= 5 920) were overrepresented at retrotransposons (LINEs and LTR elements like ERVL, ERVL-MaLR and ERVK) (z-score > 45; p-value<1×10^-3^, n = 1 000 permutations) (**Fig. 3A, 3B, Supplementary Table 7**), and these tended to gain methylation in individuals with increased BMI (**Fig. 3C**). Given that the insertion age of a TE is associated with its activity, our next step was to query TE insertion age in relation to changes in DNA methylation associated with increased BMI. To do so we applied a linear model to associate significant changes in DNA methylation due to BMI with transposable element insertion age (**Extended Data Fig.3, Supplementary Table 8**). First, we checked at the level of TE families then subfamilies. With the exception of ERVK and centr elements, no families appeared to be more likely enriched for BMI-DMCs compared to other families (**Fig. 3D**). However, some subfamilies demonstrated a strong correlation between their insertion age and the degree of DNA methylation variation associated with BMI. Notably, these included younger hominoid-specific subfamilies like HERVK elements (HERVK13-int), L1HS, L1PA14, or L1M subfamilies, as well as Alu elements (AluJr4, AluSc, AluY…) (**Fig 3E, Extended Data Fig. 4A**). This suggests that cytosines at young elements are more likely to be mis-methylated in men with increased BMI. In the human genome, it is estimated that on average, 20% of all binding events of a transcription factor fall within a transposable element ^65^. Young LINE1 elements (L1), HERVK, and Alu elements have been identified as potential long-range regulators of gene expression in human cells, playing a key role in shaping early development by influencing regulatory networks ^64,66^. Given the known role of TEs as platforms for transcription factor binding motifs, and their critical role in early development, we examined transcription factor binding motifs found at DMRs located in LINE (n= 504) and LTR (n=288) elements. Many DMRs found at these elements were associated with known regulators of early development (HOX and ONECUT factors, ZFP42, ZIC4, KLF14, ETV5). We also noticed an association with brain developmental factors (ARNT, GSX1) (p-value <0.05 and log10(enrichment) >2) (**Fig. 3F, Extended Data Fig. 4B, 4C, Supplementary Table 9**).

**Fig 3.**
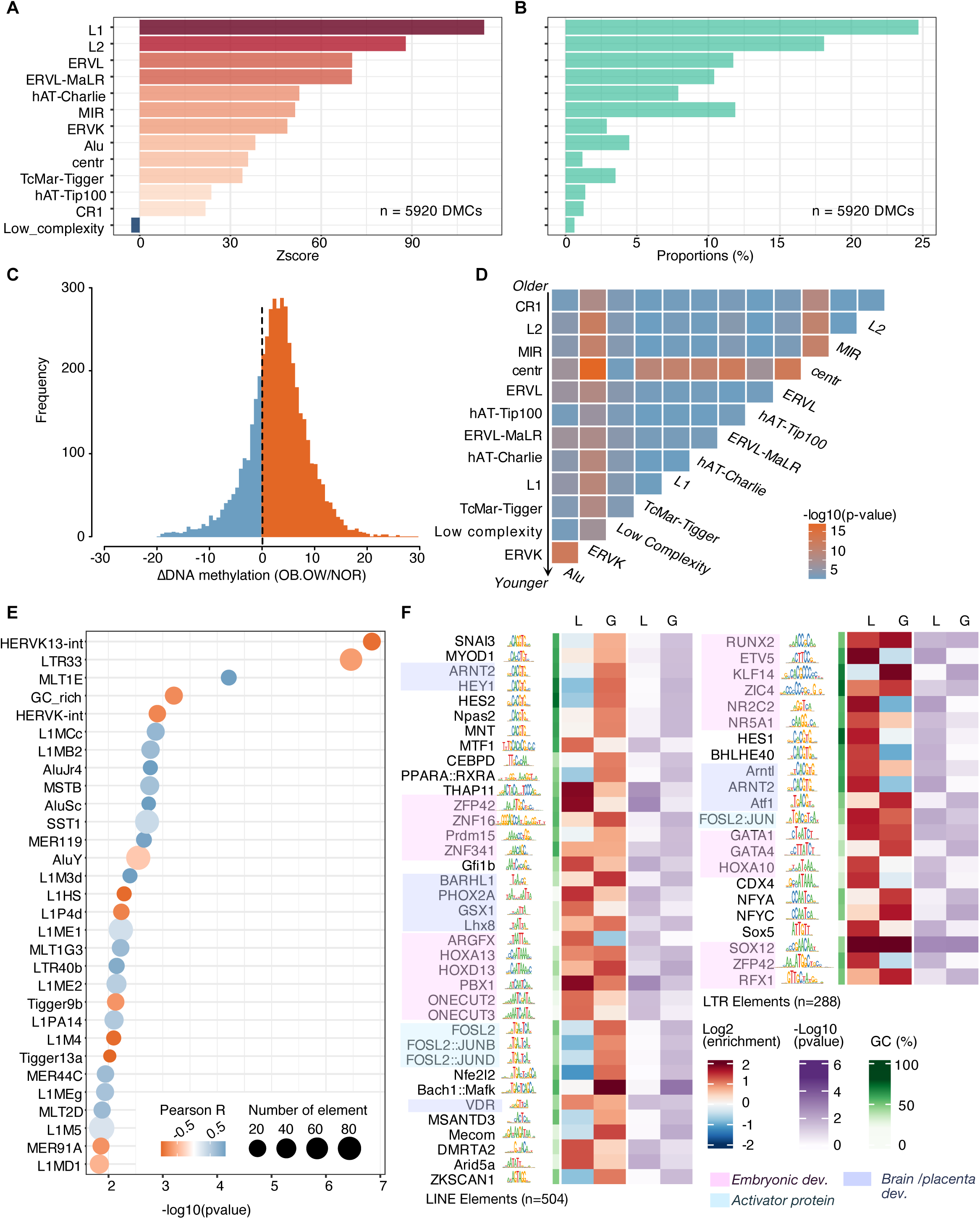
BMI may influence methylation at young retrotransposons harboring binding motifs associated with developmental transcription factors. **A**. Barplot of Zscore enrichment of BMI-DMCs on TE Families over background (n=5920). Scores were obtained using n=1000 random permutation t-test using R package RegioneR. **B**. Barplot of proportions of BMI-DMCs on TE Families in percentage (n=5920). **C**. Histogram of DNA methylation changes in OB.OW vs NOR individuals at DMCs present on Transposable elements. **D**. Matrix comparison of TE families ordered by their average age comparing their association with BMI-DMCs changes. Pairwise comparisons were made using Wilcoxon rank test. **E**. Bubble plot showing the transposable elements with most significant changes in BMI-DMC relative to the element age. Pearson R coefficient was calculated using LM model. **F**. Transcription factors motifs enrichment analysis over DMR at LINE1 and LRT Elements showing DNA methylation gains or losses (L/G). Enrichment statistics were calculated using one-sided Fischer test.

Given that both BMI and age are associated with increased inflammation, a known driver of DNA methylation alterations ^67–70^, we explored whether there was a connection between BMI, age and altered sperm methylation. To test this potential association, we applied an adjusted generalized linear regression model to measure quantitative association between sperm DNA methylation and age. Indeed, we found 97 412 DMCs associated with age. Among them, 9837 (**Supplementary Table 10**) sperm DMCs were found to be associated with both BMI and age (Shared Age/BMI DMCs) (qvalue <0.01) (**Extended Data Fig. 5A**). Like BMI-DMCs, Age-DMCs overlapped with previously identified “dynamic” CpGs ^19^ and found enriched in intergenic regions (Zscore >100), but also strongly enriched at Introns and upstream gene promoters (Zscore >200, and Zscore >100 respectively) (**Extended Data Fig. 5A, 5B**). Corresponding Age-DMRs at gene features were also associated with neural gene ontologies (p-value <2.5×10^-5^) (**Extended Data Fig. 5C, Supplementary Table 11**). This is consistent with previous work associating increased paternal age with increased risk of autism/ADHD syndrome in children ^15^. Moreover, Age-DMCs were enriched at retrotransposons (notably LINE and ERV elements, Zscore >100) (**Extended Data Fig. 5D**). Again, shared BMI/Age-DMCs demonstrated strong association with subfamilies of transposable elements that were identified at BMI-DMCs like LTR33, SST1, L1M3D elements (**Extended Data Fig. 5E**). This suggest that these subfamilies are more likely to be significantly affected by DNA methylation changes in association with both age and BMI. The consistent enrichment of both BMI-DMCs, Age-DMCs at intergenic regions that display dynamic methylation levels suggests a potential function as environment/metabolic biosensors. Indeed, after merging Age-DMCs into Age-DMRs (n= 10 329) (**Extended Data Fig. 5F**), we found some overlap between Age-DMRs with previously reported DMRs involved in published sperm epigenetic clocks ^33,44^. Moreover, some DMRs influenced by both age and BMI (n=1 304) showed overlaps with sperm epigenetic clocks DMR from *Cao et al.* and *Pilsner et al.*, (84 and 39 respectively). More saliently, 136 and 83 BMI-DMR were detected to overlap with Cao’s and Pilsner’s epiclocks, respectively. This finding highlights BMI as a potential confounding factor when considering the effects of age on the sperm methylome (**Extended Data Fig. 5F**).

### Sperm BMI-DMRs occurred in embryonic enhancers within transposable elements

Transposable elements can evade epigenetic reprogramming in the early embryo ^24,25,71^. To investigate the potential of sperm BMI-DMRs to impact early human development, we overlapped sperm BMI-DMRs with active 8 Cell-stage (8C) enhancers, a critical time point of human zygote genome activation ^47^. Enhancers were defined by the presence of Histone 3 Lysine 27 acetylation (H3K27ac), with and/or accessible chromatin marked by ATAC-seq, and showing HiC interaction with gene promoters at a minimum distance of 3kb from the nearest TSS ^47^. We identified 197 BMI-DMR enhancers that associated with 123 genes (**Fig. 4A, Supplementary Table 12**). Some of these DMR associated enhancers overlapped TEs(n=79), that were primarily ERV, LINE elements (**Fig. 4B**), aligning with previous findings showing that LTR, LINE elements to contribute to enhancer regulation in early development ^72,73^. Most DMRs localized to 8C enhancers tended to gain methylation (n=159), while 38 lost methylation (**Fig. 4A, 4C, Extended Data Fig.6D**). Given the relatively short list of genes, we detected no enriched ontologies associated with these regions. To gain insight into how DMRs associated with enhancers may influence gene expression and embryo quality, we leveraged RNA-seq data profiled in 8C human embryos over arrested-blastomeres (**Extended Data Fig. 6A, 6B, 6C**) ^46,47^. We observed that genes associated to sperm BMI-DMR at 8C enhancers were active at ZGA (**Fig. 4D**). To further characterize these overlaps between 8C Enhancers and sperm BMI-DMRs, we looked for enriched transcription factors binding motifs. Identified binding motifs included important regulators of early development, most notably embryonic and brain development factors (SOX, ASCL1, FOS:JUN, TFAP factors) (**Fig. 4E, Extended Data Fig. 6D, E, Supplementary Table 13**). Interestingly, several of these factors are involved in inflammation, or are clinical targets used to reduce inflammation like MAFG or REL ^74,75^. This may reflect an impact of BMI that is related to the chronic inflammation state of being overweight. In addition, some DMR/ZGA enhancer element associated with placental development associated factors SNAI1 and FOSL1 ^76–78^ (**Fig. 4E and Supplementary Table 13**). These associations suggests that if these DMRs are paternally transmitted and escape reprogramming, this may impact ZGA, initiation of embryogenesis and/or placentation.

**Fig 4.**
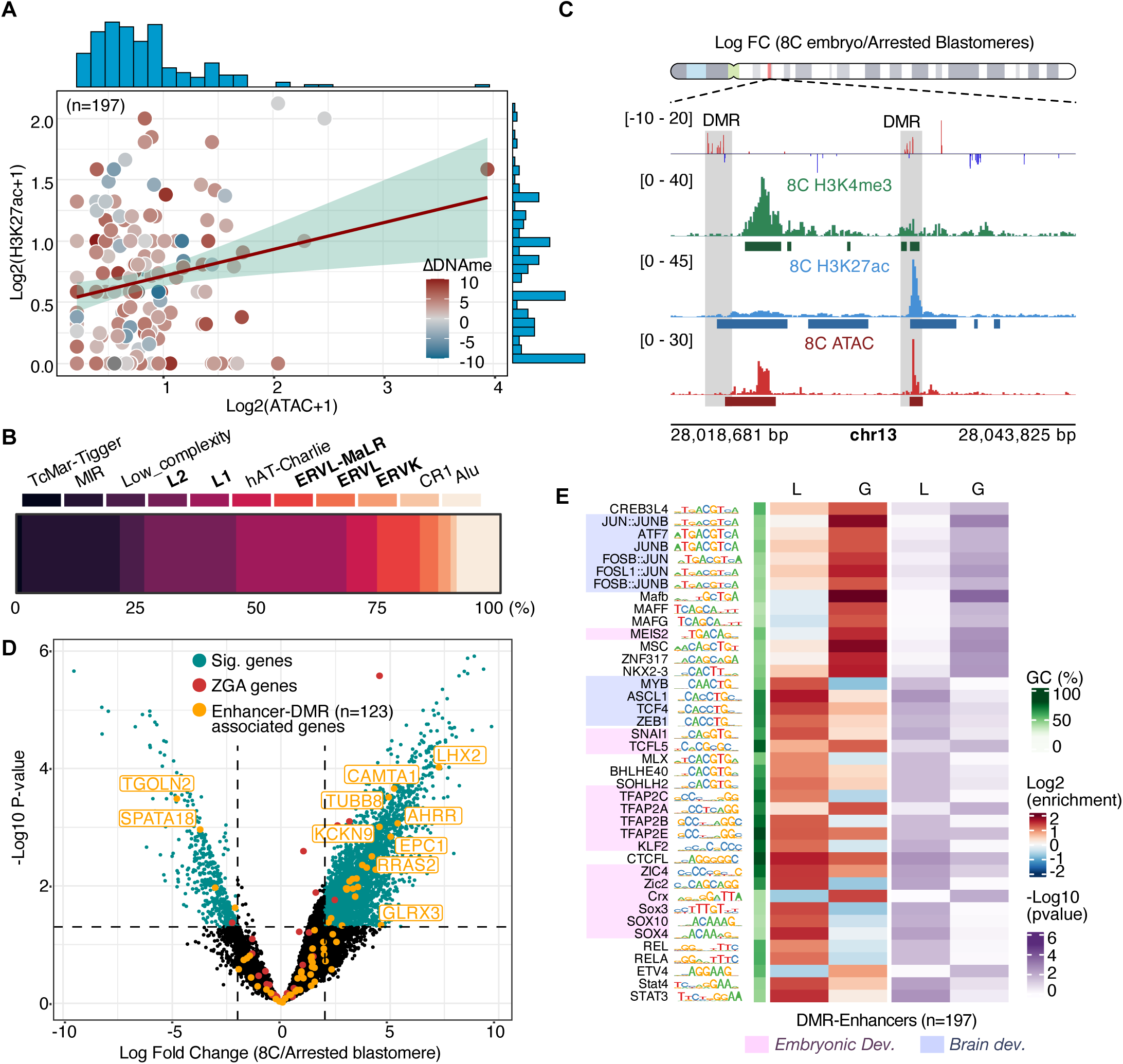
A subset of sperm DMRs overlaps with 8-cell enhancers that are expressed in 8C embryos. **A**. Scatter plot showing the level of DNA methylation, and ATAC/H3K27ac enrichment (Average log2 enrichment) at BMI-8C enhancers DMRs. **B**. Proportion of Transposable elements (n=79) found overlapping at BMI-Enhancers DMR. **C**. Signal track showing the ΔDNA methylation at DMRs overlapping with 8C enhancers marked with H3K27ac and ATAC. **D**. Volcano plots showing the expression of BMI-DMR and 8C-enhancers associated genes in normal 8C human embryos over arrested blastomeres. Thresholds were set up at FDR < 5% and Log2(FC) > 1 (two-sided t-test). Data obtained from biological duplicate (Yu et al 2022, and Wu et al 2018). **E**. Transcription factors binding motifs enrichment analysis on lost and gained (L/G) BMI-DMR 8C enhancer. Enrichment stats were calculated using one-sided Fischer test.

### A subset of BMI-DMRs overlapped with promoters of active genes in lineage specification at the blastocyst stage

It has long been established that the paternal epigenome is a key regulator of the first lineage segregation, and the development of the extra-embryonic tissues through imprinted genes ^3,79^. In pregnancies where the father is overweight, there is and increased incidence of miscarriage, placental developmental abnormalities, and obstetrical complications such as pre-eclampsia ^80^. To assess whether DMRs in sperm from men with elevated BMI could persist through development and impact trophectoderm gene expression, we queried whether 8C enhancer regions overlapped with sperm-DMRs that remain active at the blastocyst stage. Interestingly, 8C enhancers marked with sperm-BMI DMRs not only retained accessible chromatin at the ICM stage compared to the 8C stage, but even demonstrated a slight increase in H3K27ac (**Fig. 5A**). This suggests a potentially persistent effect of the altered epigenomic landscape at regions overlapping with sperm BMI-DMRs.

**Fig 5.**
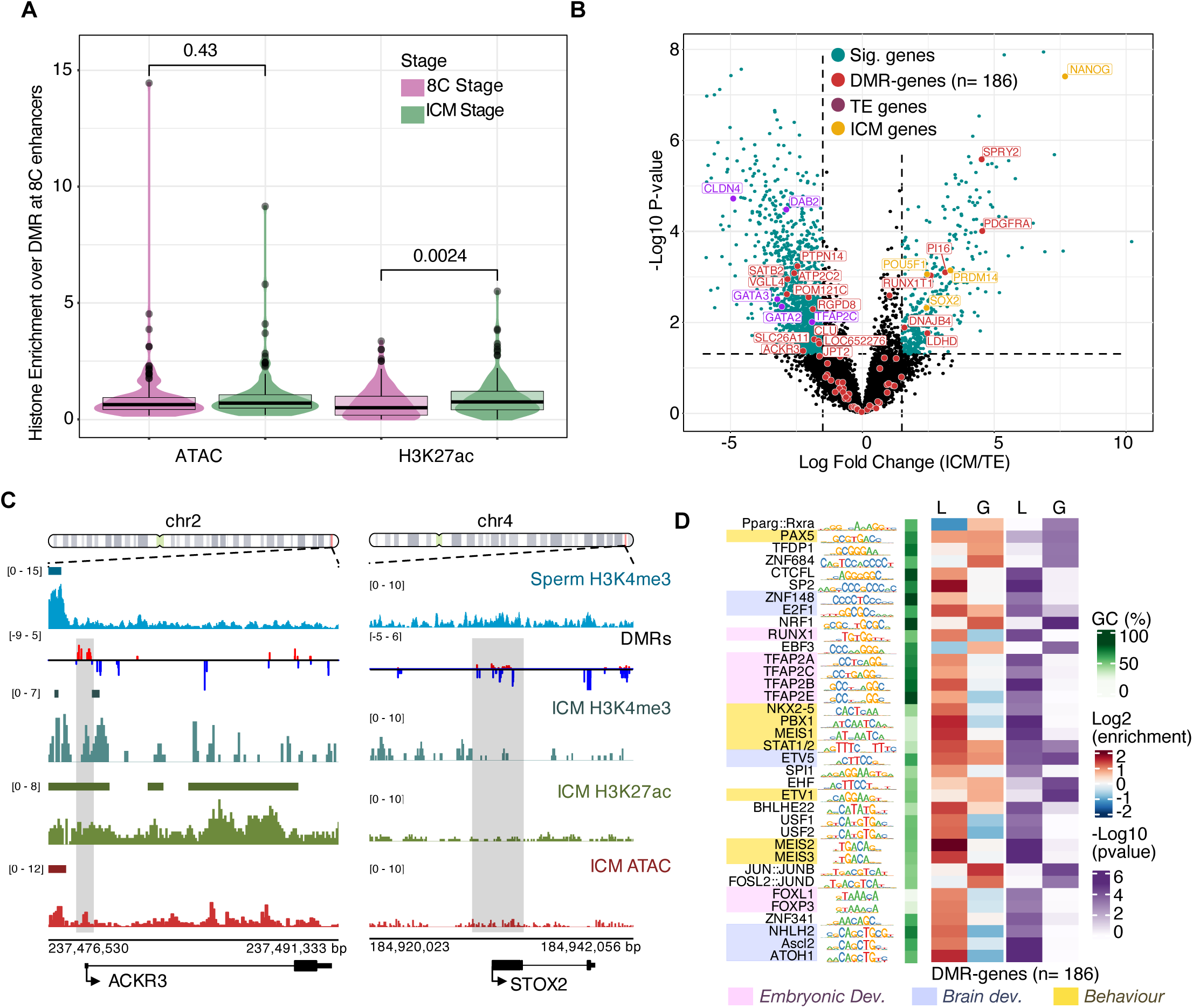
A subset of sperm DMRs overlaps with genes promoter that show a Trophectoderm biased expression in human blastocysts. **A**. Violin plots of enrichment for ATAC and H3K27ac at 8C enhancer at 8C stage and in the ICM at blastocyst stage. **B**. Volcano plots showing the expression of BMI-DMR at genes promoter and their associated expression ICM vs TE human blastocysts. Thresholds were set up at FDR < 5% and Log2(FC) > 1 (two-sided t-test). Data obtained from five biological replicate (Kai et al 2022). **C**. Signal tracks in human ICM showing the ΔDNA methylation at DMRs overlapping with gene promoters (ACKR3 inflammation related-gene and STOX2 expressed in placenta of pre-eclamptic women). **D**. Transcription factors motifs enrichment analysis on lost and gained (L/G) BMI-DMR at gene promoters Enrichment stats were calculated using one-sided Fischer test.

Next, to investigate the potential impact of sperm BMI-DMRs on cell fate decisions at the blastocyst stage, we looked at the overlap of DMRs with gene promoters. We found 477 DMRs overlapping with gene promoters and 5’UTRs (**Extended Data Fig. 7A, Supplementary Table 14**), some of which were associated with enriched H3K4me3 overlapping with ATAC peaks at the ICM stage, regardless of whether there was a gain or loss of DNA methylation level. These DMRs were associated with 186 genes. Consistent with our findings at the 8C stage, these also included inflammation genes, or genes that are clinical targets to reduce inflammation like ACKR3, S1PR3, and GAS6 (**Fig 5B, 5C, Supplementary Table 14**) ^81–83^. Interestingly, we also found regulators of spermatogenesis like TEX19, which is involved in the piRNA pathway ^84^, and STOX2 whose expression in the placenta is associated with pre-eclampsia ^85^ (**Fig 5C, Supplementary Table 14**). Despite the convergent roles in inflammation and placental development found between gene promoter overlapping with sperm BMI-DMR and gene regulated with ZGA enhancer bearing a DMR, they were not enriched gene ontology terms given the limited number of genes with DMRs involved in these processes.

We re-analyzed RNA-seq data of human early lineages, the epiblast/ICM and trophectoderm/TE ^45^ (**Extended Data Fig. 7C, 7B**) to examine the expression patterns of these genes marked by DMRs in the sperm of men with elevated BMI. We found that sperm BMI-DMR occurred at gene promoters in both ICM and TE lineages and were expressed in both lineages. A larger proportion (61%) of DMRs were associated with developing trophectoderm at the blastocyst stage (**Fig. 5B, Supplementary Table 14**). Binding motifs at these promoters revealed transcription factors associated with neurodevelopment (ASCL2, ETV5), as well as early cell fate decision transcription factors (RUNX1, TFAPs, FOX factors). In addition, TFAP factors have been shown to play an important role in regulating the emergence of the trophectoderm lineage in mice ^86^. Interestingly, these motifs were also associated with behavioral genes, which links to our disease ontology analysis (notably through MEIS factors) (**Fig. 5D, Extended Data Fig. 7D, Supplementary Table 15**). This supports the notion that BMI-sensitive DMRs at promoters could influence genes expressed in the blastocyst, and their epigenetic alteration could lead to impaired embryonic development. Moreover, the higher proportion of sperm-DMR-marked genes being expressed in the trophectoderm resonates with the known function of the paternally inherited epigenome in the development of the placenta ^3,4,79^. The placenta, a known neuroendocrine tissue, is critical in the development of the embryo’s nervous system ^87^. Hence, our data highlight possibility of a sperm-placenta-brain axis as a potential route for paternally transmitted epigenetic insults to affect early embryo development and neural development.

## Discussion

### The sperm epigenome is sensitive to elevated BMI and age

The literature highlights external factors as being influential on men’s pre-conceptional health, with elevated weight linked to impaired fertility and altered sperm methylome ^91–94^. Furthermore, the effects of paternal inheritance have been observed across generations in both human population studies and animal models ^90–93^, highlighting the consequences of men’s health on the following generation. The sperm epigenome, and DNA methylation in particular, may serve as a potential bridge between generations.

Overall, we identified over 38 000 BMI-sensitive DMCs and over 3 800 differentially BMI-sensitive DMRs, mostly located at intergenic regions in the sperm of men with elevated BMI. Supporting a role for paternal BMI influence on neurodevelopmental disorder in children we observed DMRs at genes associated with neural developmental and disease ontologies. This finding is consistent with previous studies ^94^, and further implicates sperm DNA methylation as a potential mechanism underlying the epidemiological findings that overweight and obese men are at a greater risk of having a child with a neurological disorders including autism. This association has important ramifications for prevention of childhood disease and the need to include fathers’ health as a consideration for pre-conception intervention.

Interestingly, most of the sperm BMI-DMCs/DMRs displayed a trend for hypermethylation. As we observed enrichment of DMR at promoters of genes involved in inflammation (REL, ACKR3, MAFG) (**Supplementary Table 14**), one possible explanation can be drawn from the association between obesity and low-grade chronic inflammation in the testes ^68,95–97^. Indeed, oxidative stress triggered by inflammation is known to impact the DNA methylation landscape through DNA damage repair pathways, or alteration of metabolism, and/or changes in the cell’s redox environment ^69,70^. It was proposed that some super-oxide reactive species could increase the likelihood of methylation deposition at cytosines ^98^. Furthermore, DNMTs are known to be recruited to DNA damage sites; specifically hypermethylation can be promoted via DNMT recruitment at promoters via oxidative stress ^99,100^. This is worth considering as one molecular mechanism that could explain the preponderance of DMCs bearing increased methylation.

Here we report that BMI and age may influence a common subset of DMCs, and that sperm BMI-DMR overlapped those implicated in sperm epigenetic clock models (Cao et al., 2020; Pilsner et al., 2022). Interestingly, inflammation is a common physiological responses that also occur with increasing age ^101^. This reinforces the idea that there may be a common subset of DMCs/DMRs in sperm that are responsive to multiple physiological and/or environmental conditions and diseases. Moreover, a recent study as also found that age-related change in sperm DNA methylation associate with neural ontologies, further highlighting the association between these cues ^102^. As open-source sperm DNA methylation data from men under various conditions becomes available, such a large-scale analysis using machine learning or artificial intelligence-based approaches to probe for common functional regions of the genome will be warranted.

Moreover, in a mouse model paternal obesity, oxidative stress, and inflammation have also been linked to changes in other epigenetic marks, particularly H3K4me3 ^103,104^. Our work here highlights that DMRs can occur in regions normally marked by H3K4me3. Interestingly, H3K4me3 and DNA methylation both rely on the availability of the methyl donor SAM, which is produced through the one-carbon cycle, highlighting a potentially shared metabolic dependency ^105,106^. Due to limitations in sperm number, and the use multi-omics methods not yet being optimized for sperm, we were unable to simultaneously probe other epigenetic read-outs like small non-coding RNA, chromatin accessibility or other histone modifications ^107^. In future work, a more comprehensive view of the sperm epigenome in men with elevated BMI should be pursued to explore their interconnective relationships.

### Could a “Sperm-Placenta-Brain” axis link paternal BMI to offspring disease via transposable elements?

BMI-associated DMCs significantly overlap with some families of retrotransposons (mostly LTR/ERV and LINE1). These elements have the capacity to duplicate themselves with potential deleterious effects for the host genome, and DNA methylation is a critical repressor of their activity. Notably the repression of young and potentially transcriptionally active transposable elements is of tremendous importance for spermatogenesis and fertility ^5,108,109^. Despite their potential negative effect on the genome, transposable elements are also known to act as platforms for the recruitment of transcription factors (TFs), and play the role of cis-regulatory elements, like enhancers. It is estimated that 20% of human transcription factor binding motifs are encoded in transposable element sequences ^65^. Remarkably, our binding motif enrichment analysis over BMI-DMRs at LTR and LINE1 elements indeed showed strong enrichment for transcription factors that are associated with embryonic and brain development. Additionally, we observed BMI-DMRs overlapping with transposable elements and 8-cell enhancers that are active at ZGA in human embryos up-until the blastocyst stage, a critical time point of cell fate decisions and embryo development. The consistent representation of neural pathways ontologies and neural factors motifs at DMRs over transposable elements or gene promoters is intriguing. This association between differences in DNA methylation and neural regulation has been recurrently observed in the literature both in the regulation of cell fate decisions in early development but also in men’s sperm ^110–112^. More saliently, a recent publication demonstrated potential causal association between placental DNA methylation and neuropsychiatric disorders ^113^.

In our dataset, we noted the presence of some placental genes marked by DMRs and/or regulated by ZGA enhancers overlapping with a DMR at a transposable element. Notably, STOX2 expression is detected in the placenta of pre-eclamptic women, and SNAI1, JUNB. This aligns with our observation that some sperm BMI-DMR marked gene promoters are active in the trophectoderm lineage at the blastocyst stage. Interestingly, the placenta, which emerges from the trophectoderm, is a neuroendocrine tissue that plays critical role in the regulation of embryonic development. First identified in the context of the “Placenta-heart connection” ^114^, the role of the placenta in regulating fetal brain development has since been increasingly recognized ^87^. This connection has been recently referred to as the “Placenta-Brain axis”, or “neuroplacentology” and constitutes an emerging field of research ^115,116^. Indeed, previous studies have emphasized the connection between placental dysfunction and obstetrical complications in pregnancies fathered by overweight men, as well as the increased placental serotonin output to the fetal brain in response to maternal inflammation during pregnancy ^80,117,118^. More recently, the placental origin of intergenerational effects has also been demonstrated in mice as sires with microbiome dysbiosis triggered reduce placental size and altered placental transcriptome in their offspring ^119^. Particularly pertinent for this study are our own previous findings that pregnancies sired by obese male mice result in abnormal placenta development and function ^103,104^.

Hence, we propose that DMR carried by the sperm could affect this Placenta-Brain axis. Changes in the sperm methylome could impact a key regulator of extra-embryonic tissue development and later indirectly impact embryonic brain development. From a developmental stand-point, our proposed “Sperm-Placenta-Brain” axis hypothesis is supported by the well-known unequal contribution of parental gametes, with the sperm contributing mostly to the emergence of extra-embryonic tissues in the early embryo, notably through gene imprints ^3,4^. The trend in mice and humans suggest that they are fewer regions that are paternally imprinted than maternally imprinted ^120,121^. Yet, previous work has shown that there exist transient and tissue specific imprinted regions that have life-long consequence for the offspring ^122,123^. The mechanism by which canonical and transient imprints evade epigenetic reprogramming in early mammalian development is well established, and usually involving targeted DNA methylation maintenance via KRAB-ZFPs ^2,124–126^ or an epigenetic switch ^127–130^.

How sperm DMCs/DMRs are transmitted and maintained in early embryogenesis has not been mechanistically explored. However, we propose the following hypotheses: Firstly, it has been shown that in some instances, the formation of maternal imprints can arise from the expression of LTR transposable elements (termed LTR-initiated transcriptional units; LITs) that promote the deposition of DNA methylation in sequences located downstream from these LITs ^71,131^. Indeed we observed sperm DMRs and DMCs occurring at LTR elements some of which evade reprogramming ^7,25^. Whether such events also occur in the sperm methylome remains to be investigated.

Secondly, reconstructive inheritance through an epigenetic change that triggers changes in downstream signaling represents an interesting possibility for the transmission of epimutation from sperm to offspring. For example, inheritance of transcription factors bound to the sperm chromatin and sperm chromatin genome organization through CTCF have been shown to be passed down from father to their offspring and influence development ^132,133^. This resonates with our observations of large enrichment of TFs binding motifs at sperm-DMRs, including CTCFL, that we observed in DMR-marked 8C enhancers. Interestingly, a mouse model of paternal inheritance of epigenetically edited CGI suggest that DNA methylation could be both replicatively (for highly methylated CGI) and reconstructively (for lowly methylated CGI) inherited from father to offspring ^134^.

To validate the significance of our findings and those of other associative studies of epigenetic inheritance in humans, future studies should focus on the mechanisms that allow for the transmission of such DMCs/DMRs from the sperm to the embryo. These studies require paired parental tissues and embryonic tissues to tease out the epigenetic contributions in an allele aware analysis- a daunting task given the extreme limited availability of human embryonic material. If such reproductive material is made available the application of multi-omics low-input technologies, long read sequencing and epigenome editing tools will allow for the discovery and validation of epigenomic regions that may be implicated in disease transmission. Nevertheless, our work underscores the critical need to consider men’s preconceptional health, particularly in the context of obesity, as a determinant of offspring development and health. By revealing the extensive epigenetic alterations in the sperm of men with elevated BMI, our findings provide a foundational framework for advancing our understanding on paternal factors that may influence early embryonic development and the long-term health outcomes in subsequent generations. This emphasizes the importance of integrating paternal health into reproductive and public health strategies to optimize both paternal and offspring well-being.

## Acknowledgment

We thank the many members of the Kimmins lab for their support, comments and discussion. The funders played no role in study design, data collection and interpretation, publication or the redaction of the manuscript. M.S. was supported by a post-doctoral grant from the FRQS (Fonds de recherche du Quebec en Santé #BF15-352882). This research was supported by Canadian Institutes of Health Research Grants (#341268) to S.K., R.L., H.W., C.L. and S.M., (#180414) to S.K., C.L. and S.M. and (#148425 and #191775) to J.T., CIHR DOHaD Team grant (#358654) to S.K., J.T, H.W., C.L. and S.M NRC New Beginnings Initiative grant (#INBR5-001001-1) to X.S., and S.K.

## Author contributions

M.S., X.S., J.T., and S.K. conceived the study. Sample collection and participants physiological parameters determination were performed by A.J.M., S.M., C.L., H.W. MethylC capture was performed by, C.L., R.L. and M.C.D. Method development of MCC-seq was made by D.C., T.P., G.B., and J.T. Bioinformatic analysis were conducted by X.S. and M.S. Then, M.S., S.K. X.J., and J.T interpreted the data and wrote the manuscript. All authors reviewed and approved the manuscript.

## Data availability

The data supporting the finds can be found in the main text, supplementary figures and tables. Data for BMI-DMCs/DMRs and Age-DMCs/DMRs are currently being uploaded into EGA, and will be made available to reviewers upon request. No original code was written for bioinformatic analysis and visualization displayed in this manuscript; all information can be found in the listed packages.

---

**Extended Data Fig. 1.**
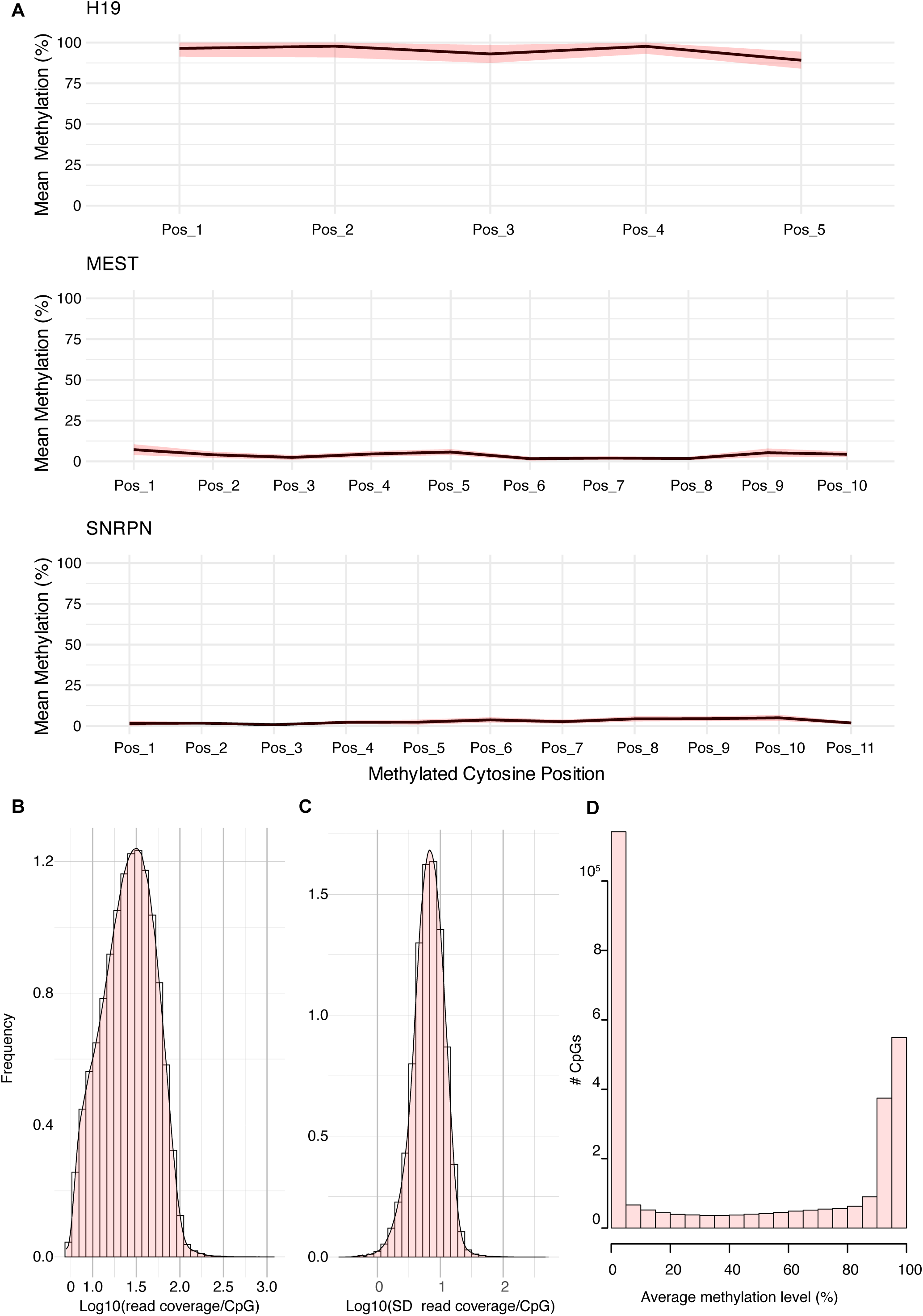
Sperm gDNA purity assessment and MCC-seq sequencing Metrics. **A**. Line plots showing the methylation at paternal (H19) and maternal (MEST, SNRPN) imprinted genes for targeted cytosines position via pyrosequencing. Black line is average methylation with standard deviation (red ribbon) across participants. Somatic contamination of sperm DNA preparation is detected with methylation levels below 90% at paternal imprints and over 10% at maternal imprints. **B**. Density plot showing the frequency of read coverage per CpG (log10) for MCC-sequencing. The coverage of CpGs was 30X in average. **C**. Density plot showing standard deviation of read coverage per CpG (log10) for MCC-sequencing. **D**. Histogram of the distribution of average methylation level (%) in the sperm.

**Extended Data Fig. 2.**
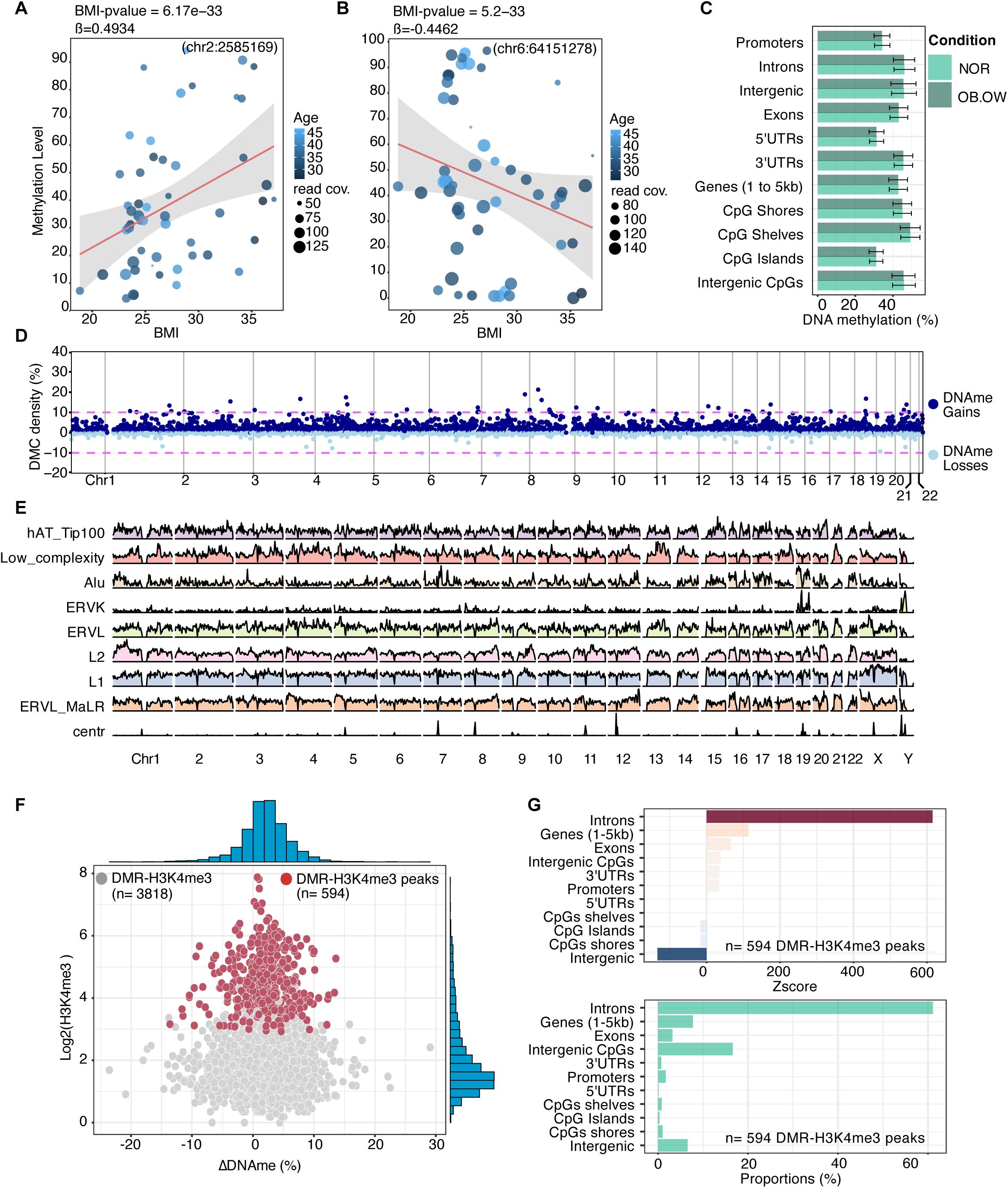
Differential CpG detection associated with BMI across sperm methylome. **A, B**. Scatter plots showing methylation levels gains and losses for all participants at single CpG location significantly associated with increased BMI. **C**. Barplot of average DNA methylation levels measured across NOR and joint OB.OW individuals at genomic features. **D**. Hotspot density analysis of significant DMCs with DNA methylation (DNAme) gains and losses over the human genome (hg19). **E**. Density distribution of transposable element families in the human genome (hg19). **F**. Scatter plot of enrichment of Sperm H3K4me3 (Lambrot, 2021) with ΔDNA methylation levels (OB.OW/NOR) at BMI-DMRs. Red dots represent the overlap between a DMR and a H3K4me3 significant peak (n=594). The blue histograms represent the density distribution of the overlaps alongside the H3K4me3 enrichment axis and ΔDNA methylation axis. **G**. Barplot of Zscore enrichment (top) of BMI-DMRs overlapping with sperm H3K4me3 peaks on key genomic features over background (n=594). Scores were obtained using n=1000 random permutation t-test using R package RegioneR. Barplot of proportions (bottom) of BMI-DMRs overlapping with H3K4me3 (n=594) on key genomic features shown in percentage.

**Extended Data Fig. 3.**
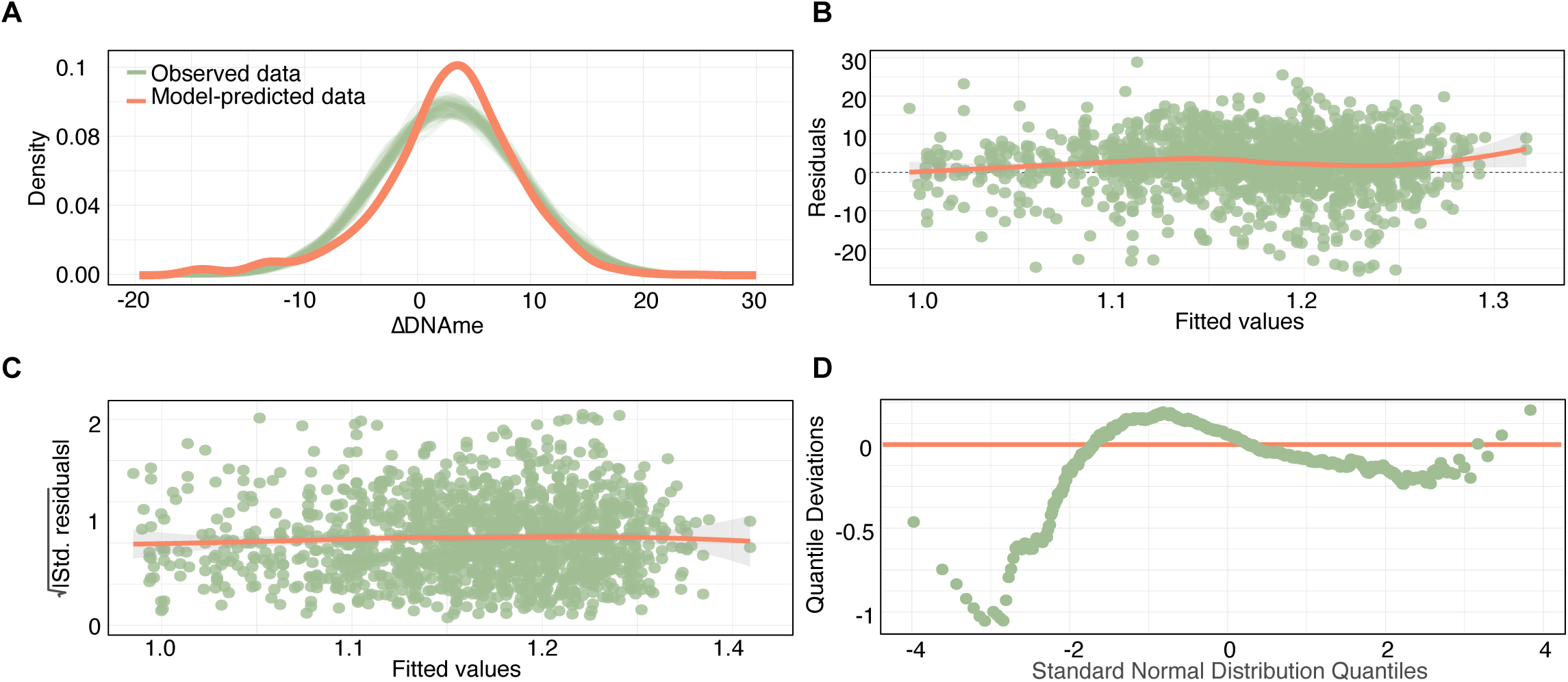
Linear model parameter for significance of BMI-DMC changes associated with element Age. **A**. Predictive check for observed changes in DNA methylation changes in BMI-DMC in association with TEs’ age vs model-predicted data. **B**. Scatter plot showing the linearity of Residuals over fitted values. Effective modelling assumes homogeneity of predictors over the outcome. **C**. Scatter plot showing the homogeneity of variance of fitted values. The models assume homoscedasticity. **D**. Scatter plot showing the normality of residuals the assess if the residuals of the regression model are normally distributed.

**Extended Data Fig. 4.**
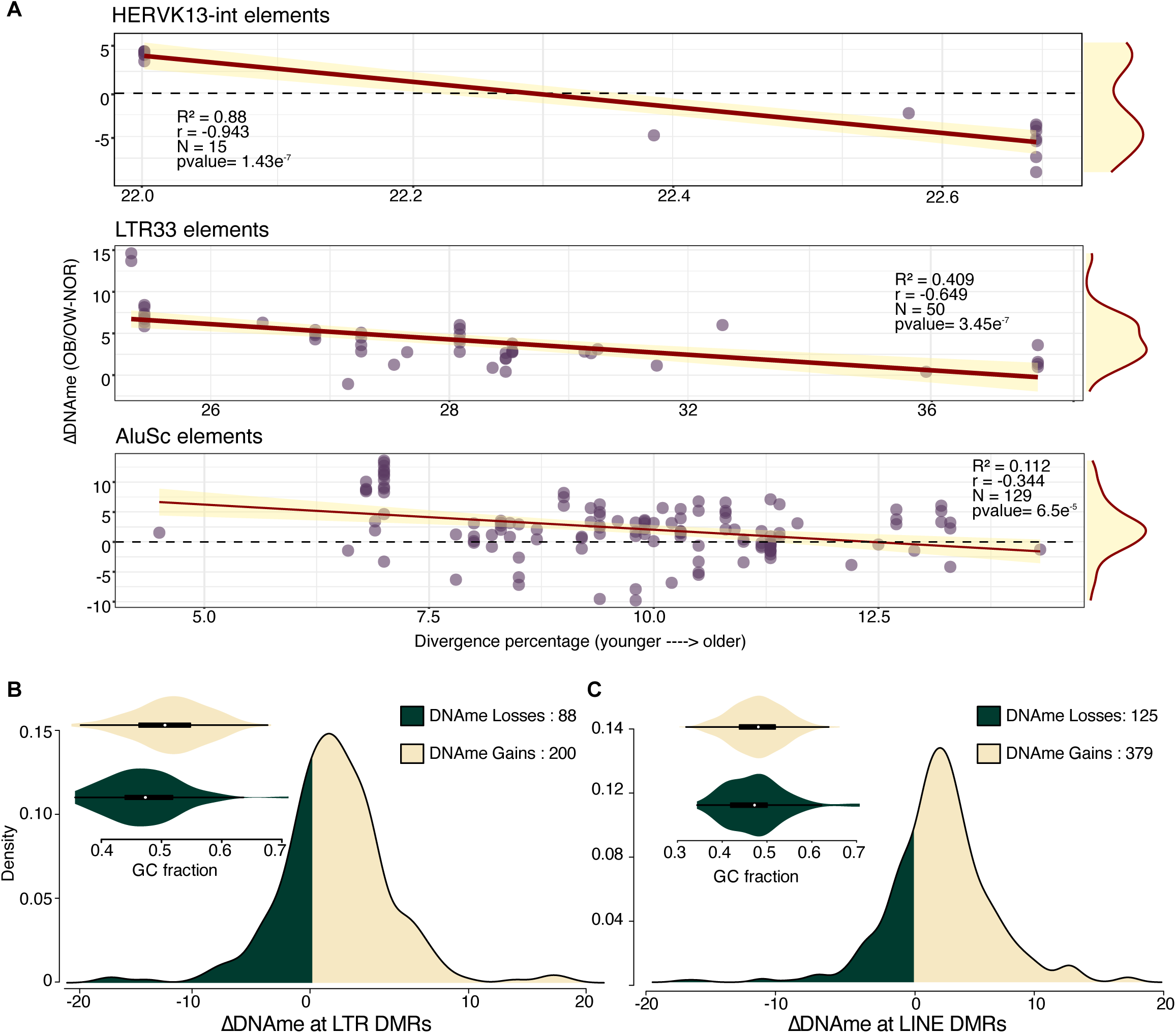
BMI-DMR preferentially occur at younger transposable element. **A**. Scatter plot showing DNA methylation changes at BMI-DMCs found at each element from specific TE subfamily according to their age. Statistics are from LM model (R2: correlation coefficient, r: pearson coefficient, N= number of elements bear BMI-DMCs from this family, p-value). **B**. Violin plot showing the distribution of GC fraction across BMI-DMRs at LTR elements with DNA methylation gains or losses. **C**. Violin plot showing the distribution of GC fraction across BMI-DMRs at LINE elements with DNA methylation gains or losses.

**Extended Data Fig. 5.**
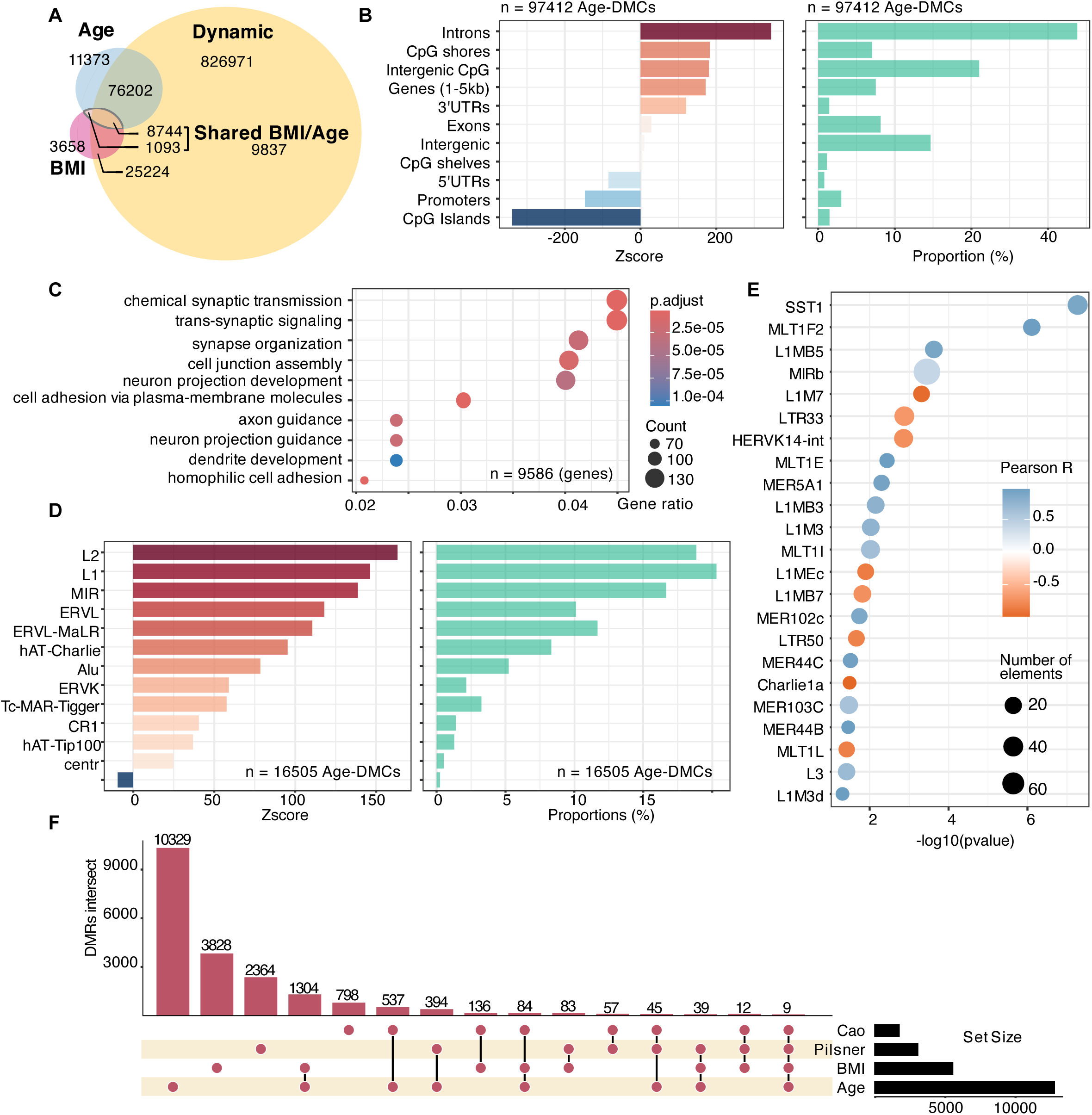
BMI-DMCs are found to overlap with Age-sensitive DMCs and also sperm epigenetic clocks. **A**. Venn diagram showing the overlap of Sperm BMI-DMCs with Age sensitive DMCs and previously defined dynamic CpGs (Chan, 2019). **B**. (left) Barplot of Zscore enrichment of Age-DMCs on key genomic features over background (n= 97412). Scores were obtained using n=1000 random permutation t-test using R package RegioneR and (right) Barplot of proportions of Age-DMCs on key genomic features in percentage (n= 97412). **C.** Bubble plot of Gene ontologies enrichment analysis (R package ClusterProfiler) for Age-DMRs located at gene promoters, introns or exons. Statistics are t-test for enrichment over background with BH correction. DMRs were defined using 500bp tile with 3 or more DMCs. **D.** (left) Barplot of Zscore enrichment of Age-DMCs on TE Families (n= 16505) over background. Scores were obtained using n=1000 random permutation t-test using R package RegioneR and (right) Barplot of proportions of Age-DMCs on TE Families (n= 16505). **E**. Bubble plot showing the transposable elements with most significant changes at shared BMI/Age-DMC relative to the element age. Pearson R coefficient was calculated using LM model. **F**. Upset plot showing overlap of DMRs sensitive to BMI, Age, Shared BMI/Age and DMRs found in Sperm epigenetic clocks (Pilsner 2022, Cao 2020).

**Extended Data Fig. 6.**
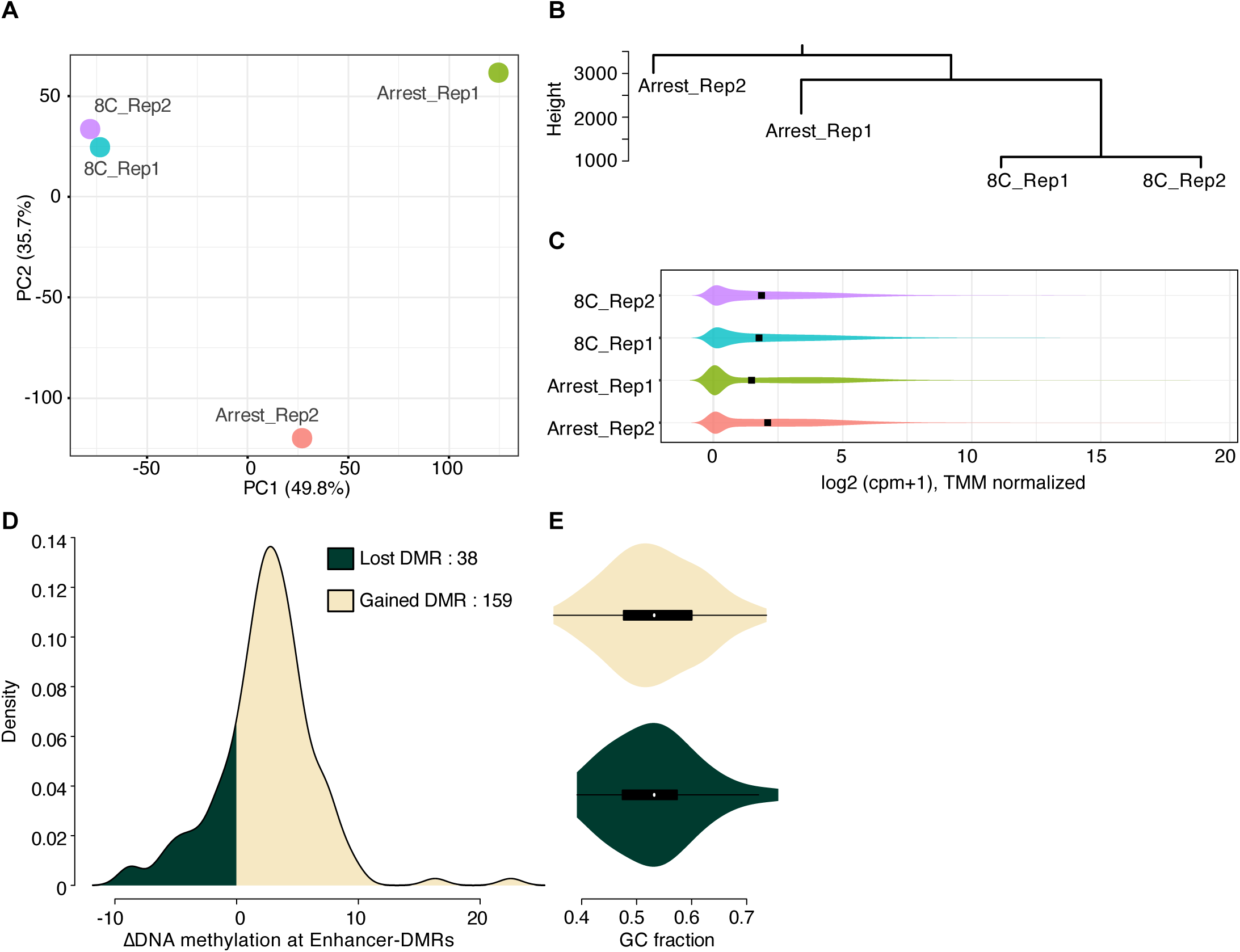
Transcriptomic analysis of 8 Cell-stage /Arrested blastomeres in relation to sperm DMRs. **A**. PCA analysis of 8C human embryo and arrested blastomeres (Xia et al 2019 and Wu et al 2018). **B**. Cluster dendrogram of 8C human embryo and arrested blastomeres (Xia et al 2019 and Wu et al 2018). **C**. Distribution of genes expression in 8C embryo and arrested blastomere displayed as their log2 level of expression. Expression was measured in CPM+1 with TMM normalization (Xia et al 2019 and Wu et al 2018). **D**. DNA methylation changes in at BMI-DMR overlapping with 8C Enhancers. **E**. Violin plot showing the distribution of GC fraction across hypo/hyper BMI-DMR overlapping with 8C Enhancers.

**Extended Data Fig. 7.**
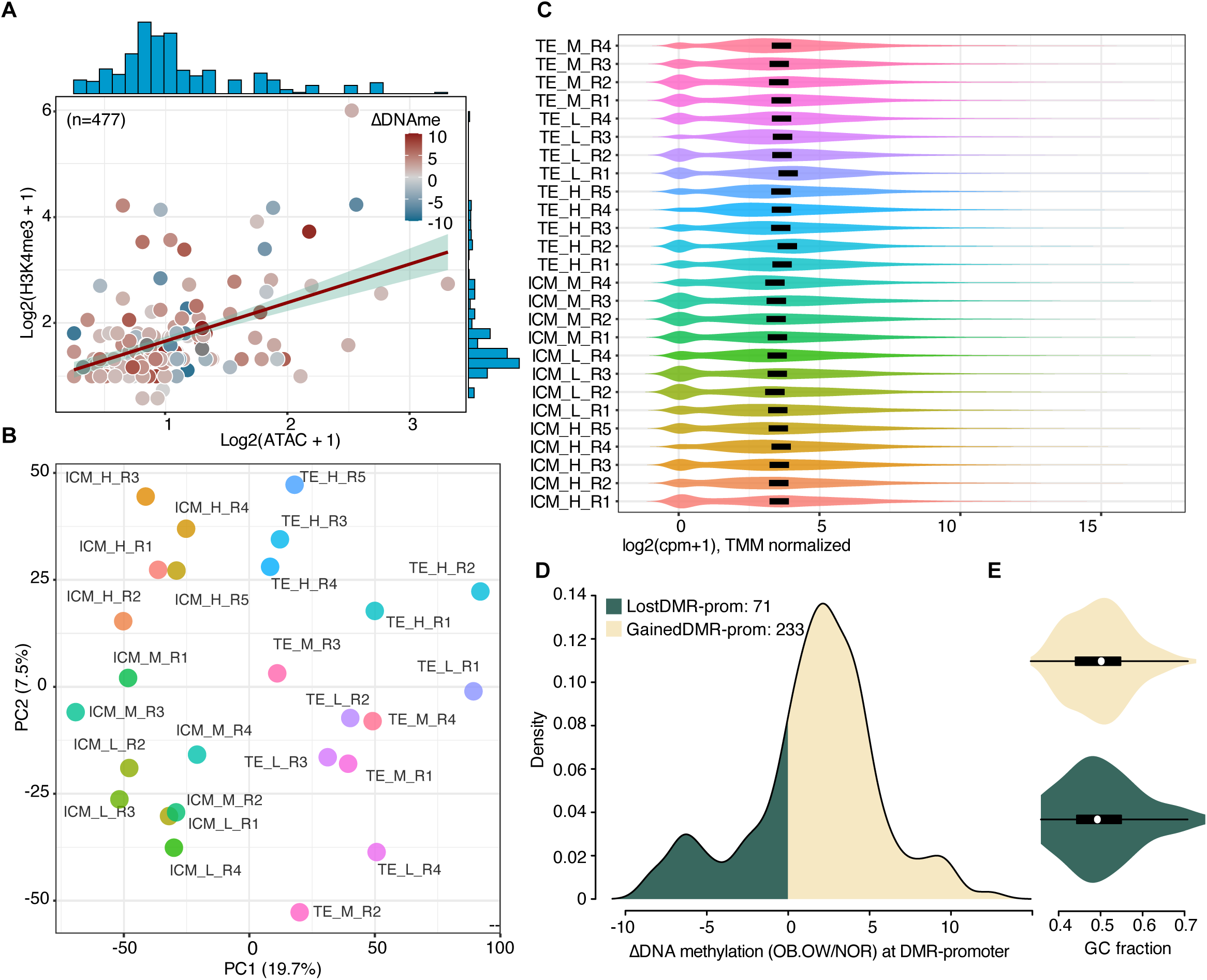
Transcriptomic analysis of ICM and Trophectoderm lineages for human blastocysts in relation to sperm DMR at gene promoters. **A**. Scatter plot showing the level in DNA methylation, and ATAC/H3K4me3 enrichment (Average log2 enrichment) at BMI-DMRs at gene promoters. **B**. PCA analysis of human blastocysts (Kai et al 2022). Replicates are grouped by pregnancy expected rates (High, mid, low). For main Fig and subsequent analysis, only High rated embryos are used. **C**. Distribution of genes expression in Human blastocysts lineages (TE and ICM) displayed as their log2 level of expression. Expression was measured in CPM+1 with TMM normalization (Kai et al 2022). **D**. DNA methylation changes in at BMI-DMR overlapping with gene promoters. **E**. Violin plot showing the distribution of GC fraction across hypo/hyper BMI-DMR overlapping with gene promoters.

